# PRISM: A High-Throughput Simulation Infrastructure for CADD Agents

**DOI:** 10.64898/2026.04.02.716083

**Authors:** Zhaoqi Shi, Xufan Gao, Mingyu Xu, Xuanyi Zhu, Peng Wang, Yuxuan Yang, Zaixing Yang, Ruhong Zhou

## Abstract

Despite rapid progress in AI agents for computer-aided drug design (CADD), protein-ligand simulation workflows remain fragmented across disparate tools, creating a major bottleneck for scalable candidate evaluation. Here, we present **PRISM** (**P**rotein-**R**eceptor **I**nteraction Simulation **M**odeler), a Python platform built on GROMACS that unifies ligand parameterization across multiple force fields, automated system construction, enhanced sampling, multi-tier binding free energy estimation, and trajectory analysis within a single workflow. Through the Model Context Protocol (MCP), PRISM further serves as the computational infrastructure for **CADD-Agent**, an expert-workflow-driven AI agent designed to orchestrate hierarchical drug screening pipelines. As a pilot application, we applied PRISM to riboflavin synthase and demonstrated end-to-end automation from candidate library assembly to binding pocket characterization, identifying a potential allosteric inhibition site at the oligomerization interface. Together, these results establish PRISM as a high-throughput simulation infrastructure for agent-enabled CADD.

## 1. INTRODUCTION

Molecular simulation has become an essential component of protein-ligand research, providing atomistic insights into binding, stability, and conformational dynamics that complements docking and other structure-based approaches. Beyond mechanistic interpretation, simulation outputs are increasingly being used in downstream data-driven modeling and molecular design. As a result, protein-ligand simulation workflows are playing an expanding role in computer-aided drug design (CADD). However, preparing simulation-ready systems in a manner that is scalable, reproducible, and compatible with downstream analysis remains labor-intensive and fragmented across multiple tools.

Several platforms have helped lower this barrier. However, important limitations remain. CHARMM-GUI^1,2^ provides accessible web-based preparation of biomolecular systems, yet its interactive submission model is less well suited for repeated or high-throughput protein-ligand workflows. OpenMMDL^3^ offers an integrated setup-simulation-analysis workflow within the OpenMM ecosystem, whereas CHAPERONg^4^ emphasizes automated GROMACS simulations together with extensive post-simulation analysis and support for biased workflows. Nevertheless, these tools do not fully bridge the gap between routine system preparation and a reproducible, end-to-end, GROMACS-native workflow that also supports flexible ligand parameterization and interpretable downstream analysis. In particular, CHAPERONg still relies on external ligand-parameter generation services such as LigParGen^5^, underscoring the continued fragmentation of many existing workflows.

This fragmentation becomes more restrictive in practical discovery settings, where users often need to evaluate large ligand sets, compare parameterization strategies, rerun simulations iteratively, and connect structural trajectories with energetic or interaction-level analyses. Although script-based automation can reduce manual effort, stage-specific pipelines are often brittle when workflows must adapt to changing task order, parameter choices, or downstream decision criteria. What is therefore needed is not merely greater automation, but a more coherent computational for end-to-end execution, reproducibility, and iterative screening.

Recent progress in scientific AI agents suggests a complementary strategy in coordinating such complex workflows. Systems such as ChemLint^6^ and Biomni^7^ have shown that large-language-model-based agents can decompose multistep scientific tasks and invoke external tools in chemistry-related settings^8,9^, while recent reviews have highlighted the importance of domain-specific, tool-grounded, and benchmarkable agent systems for real scientific applications^10^. In molecular simulation, however, the value of an agent layer depends on the availability of a robust computational backend. Rather than replacing established simulation workflows, agents are most useful when they can orchestrate setup, simulation, and analysis on top of a reproducible and well-integrated infrastructure.

Here we present **PRISM** (**P**rotein-**R**eceptor **I**nteraction **S**imulation **M**odeler), a Python platform with GROMACS interfaces for integrated, high-throughput protein-ligand simulation workflows. PRISM is designed to improve end-to-end workflow coherence by unifying automated system preparation, flexible ligand parameterization, standardized simulation configuration, and downstream analysis within a single environment. It further supports advanced tasks including MM/PB(GB)SA analysis through gmx_MMPBSA^11^, PMF-related workflows, and interaction-focused interpretation of simulation results. Relative to OpenMMDL and CHAPERONg, PRISM is intended to fill a complementary niche defined by tighter GROMACS-centered integration, stronger emphasis on reproducibility, and greater readiness for iterative and high-throughput studies. Built on this simulation infrastructure, the PRISM-CADD-Agent layer serves as a workflow-orchestration extension rather than the primary methodological claim, enabling more natural coordination across multistep CADD pipelines.

## 2. IMPLEMENTATION

### 2.1 Unified Ligand Force Field Generation

PRISM provides a unified interface to multiple ligand force-field generation pathways. Users can select one through a single command-line flag: GAFF^12^ and GAFF2 via AmberTools^13^ (Antechamber atom typing, AM1-BCC^14^ charges, tleap parameterization, ACPYPE conversion to GROMACS format); OpenFF via the OpenFF toolkit^15^ and SMIRNOFF direct chemical perception with Interchange-based format conversion; CGenFF through parsing of pre-generated CGenFF^16^ stream files; OPLS-AA via the LigParGen server^5,17,18^; and MMFF, MATCH^19^, and a hybrid scheme via the SwissParam web service. Beyond the default AM1-BCC charges, an optional Gaussian-RESP^20^ module generates electrostatic potential calculations at the HF/6-31G* or B3LYP/6-31G*^21^ level of theory and replaces AM1-BCC charges with RESP-fitted values for production-quality electrostatics. Regardless of the chosen pathway, all outputs conform to a standardized format (GRO coordinates, ITP topology, atom type definitions, and position restraint files), ensuring that downstream modules operate identically across all parameterization routes.

### 2.2 Automated Protein-Ligand System Construction

PRISM automates the complete system assembly pipeline (Figure 1). The input protein structure is validated and repaired using PDBFixer^22^ (missing heavy atoms, incomplete side chains, duplicated alternate conformers), with optional PROPKA^23^-based pKa prediction for pH-dependent protonation state assignment. GROMACS pdb2gmx generates the protein topology under a user-selected protein force field (auto-detected from the local GROMACS installation). Ligand coordinates are merged with the protein, the combined topology is assembled with appropriate cross-references, and the system is solvated in a configurable periodic box (cubic, truncated octahedral, or dodecahedral; default edge distance: 1.5 nm) with charge neutralization and ionic strength adjustment (default: 0.15 M NaCl).

**Figure 1.**
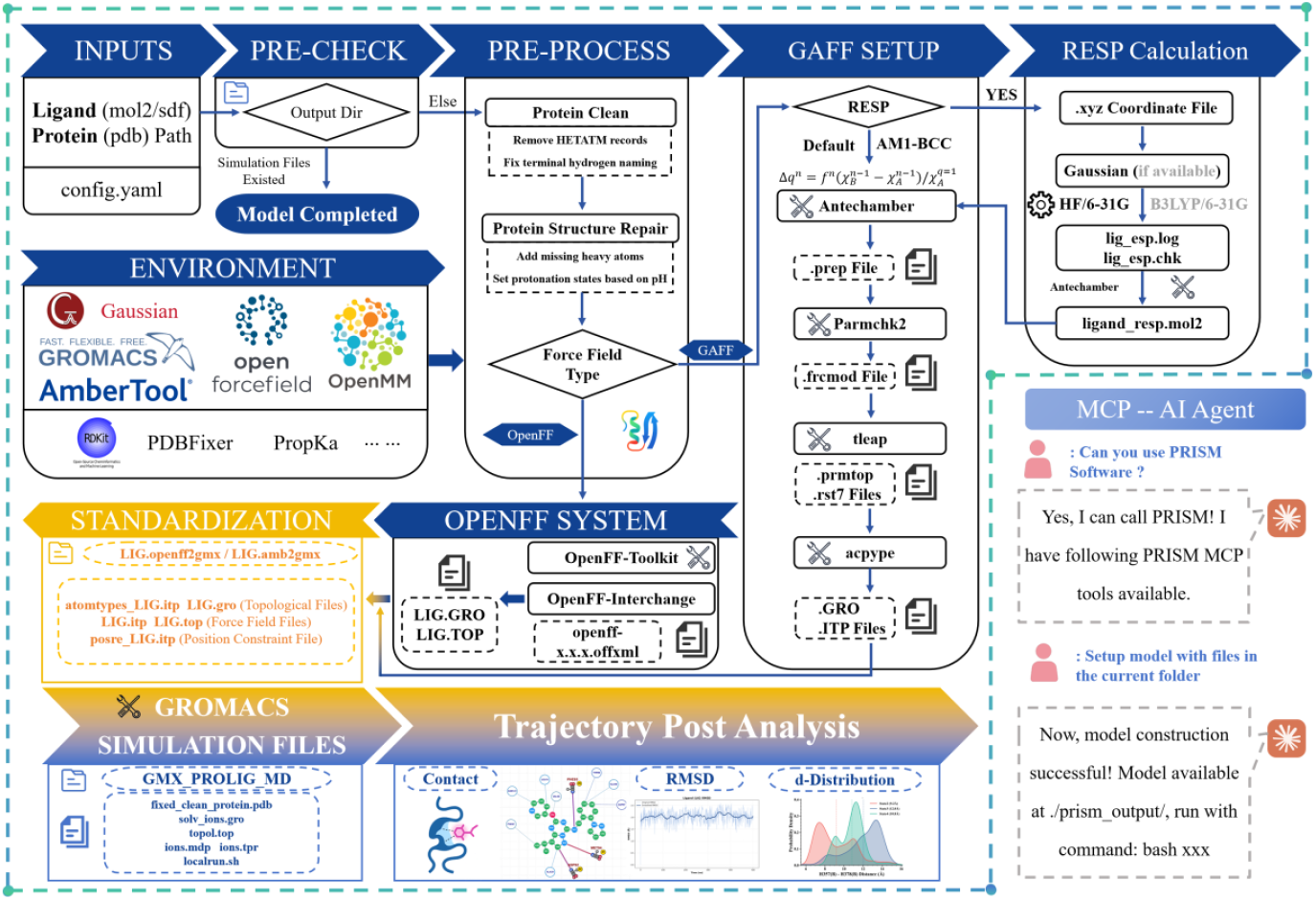
Architecture of PRISM. The platform integrates five sequential stages: input validation, protein pre-processing (PDBFixer repair, PROPKA protonation), multi-pathway ligand parameterization (GAFF/GAFF2 via AmberTools, OpenFF via Interchange, with optional Gaussian RESP charges), standardized output generation (GRO, ITP, force field and position restraint files), and trajectory post-analysis. The right panel shows the MCP interface enabling agent-driven simulation setup.

**Figure 2.**
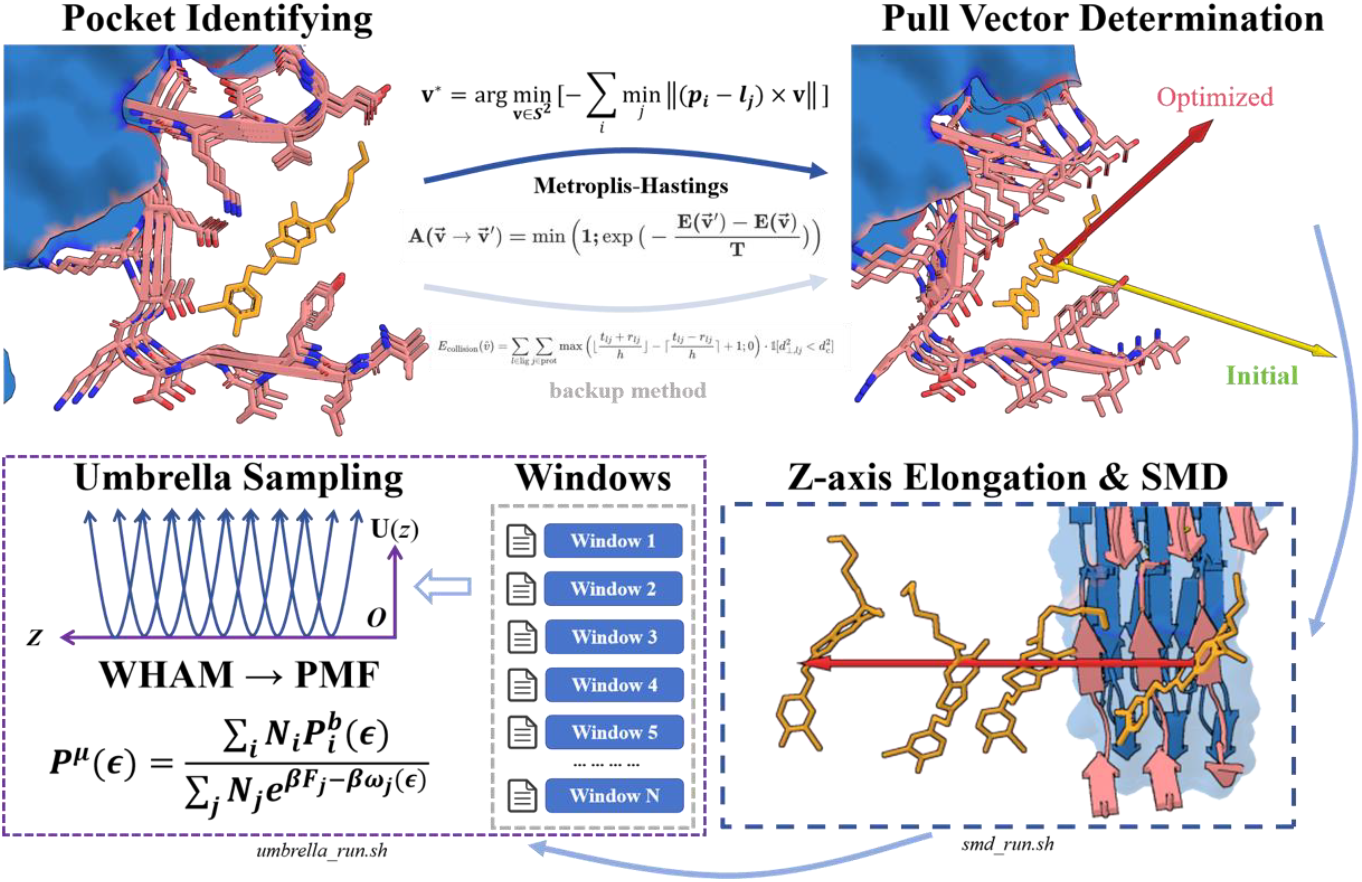
Automated PMF setup procedure. The pulling direction is optimized by minimizing a steric hindrance objective on the unit sphere S^2^ via Metropolis-Hastings sampling with simulated annealing (upper panels; initial direction in yellow, optimized in red). The complex is then rotated to align the optimal vector with the z-axis, the box is elongated, and steered MD generates a continuous unbinding trajectory from which umbrella sampling windows are extracted at uniform spacing (lower panels). WHAM reconstructs the free energy profile.

**Figure 3.**
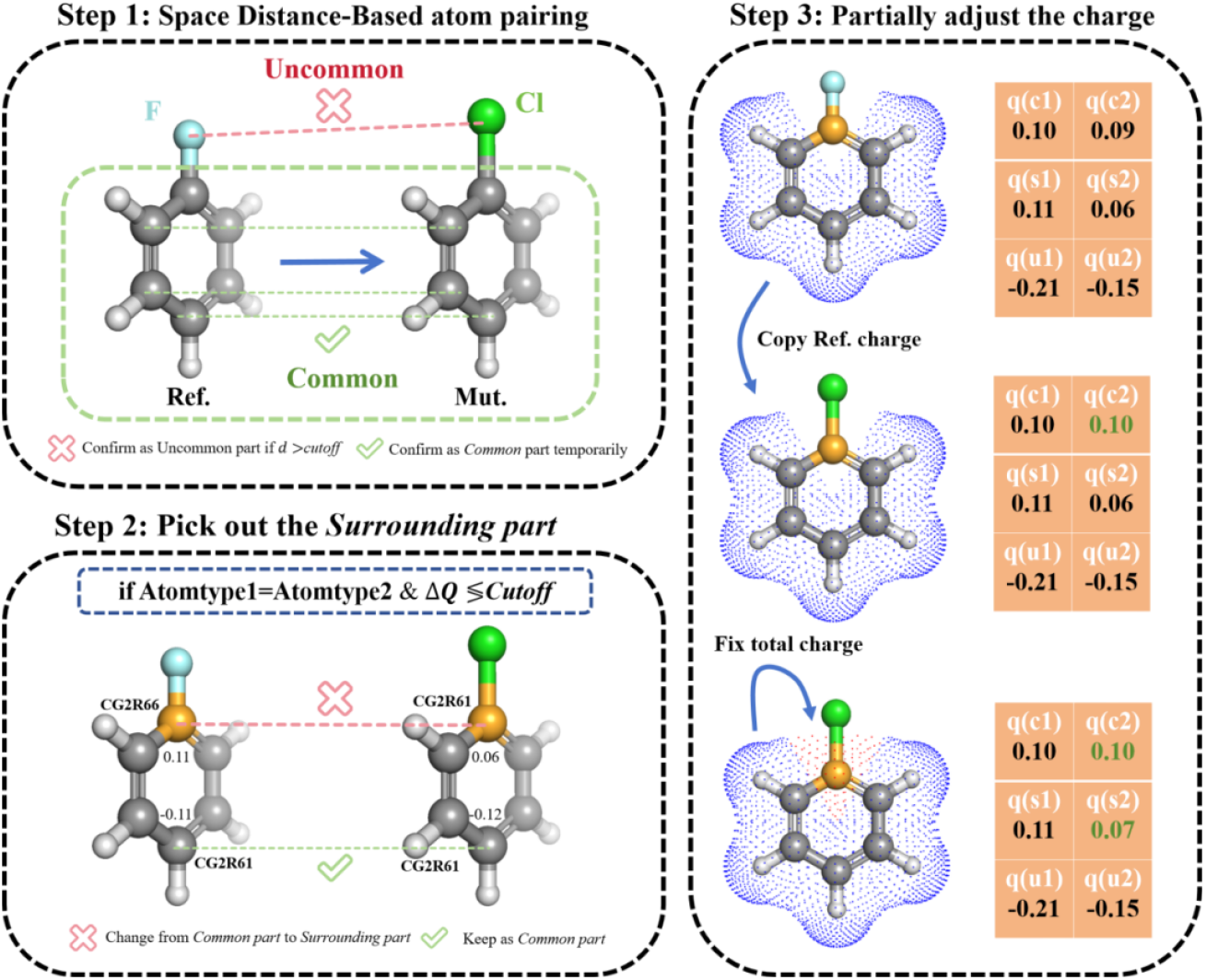
Algorithm of PRISM-FEbuilder. Schematic illustration of distance-based atom mapping and charge handling strategies for FEP system construction. (Left) The mapping algorithm identifies the shared core between reference and transformed ligands using distance criteria (0.6 Å cutoff) and element type matching, classifying atoms as common (shared scaffold), transformed (state-specific), or surrounding (position-matched but parameter-divergent). (Right) Charge assignment handles electrostatic differences across the perturbation through three strategies: reference-state preservation, mutant-state preservation, or arithmetic averaging for common atoms with minor charge differences. The single-topology GROMACS format encodes state-specific parameters via typeB/chargeB columns, enabling alchemical transformations without the parameter merging overhead required in dual-topology approaches.

**Figure 4.**
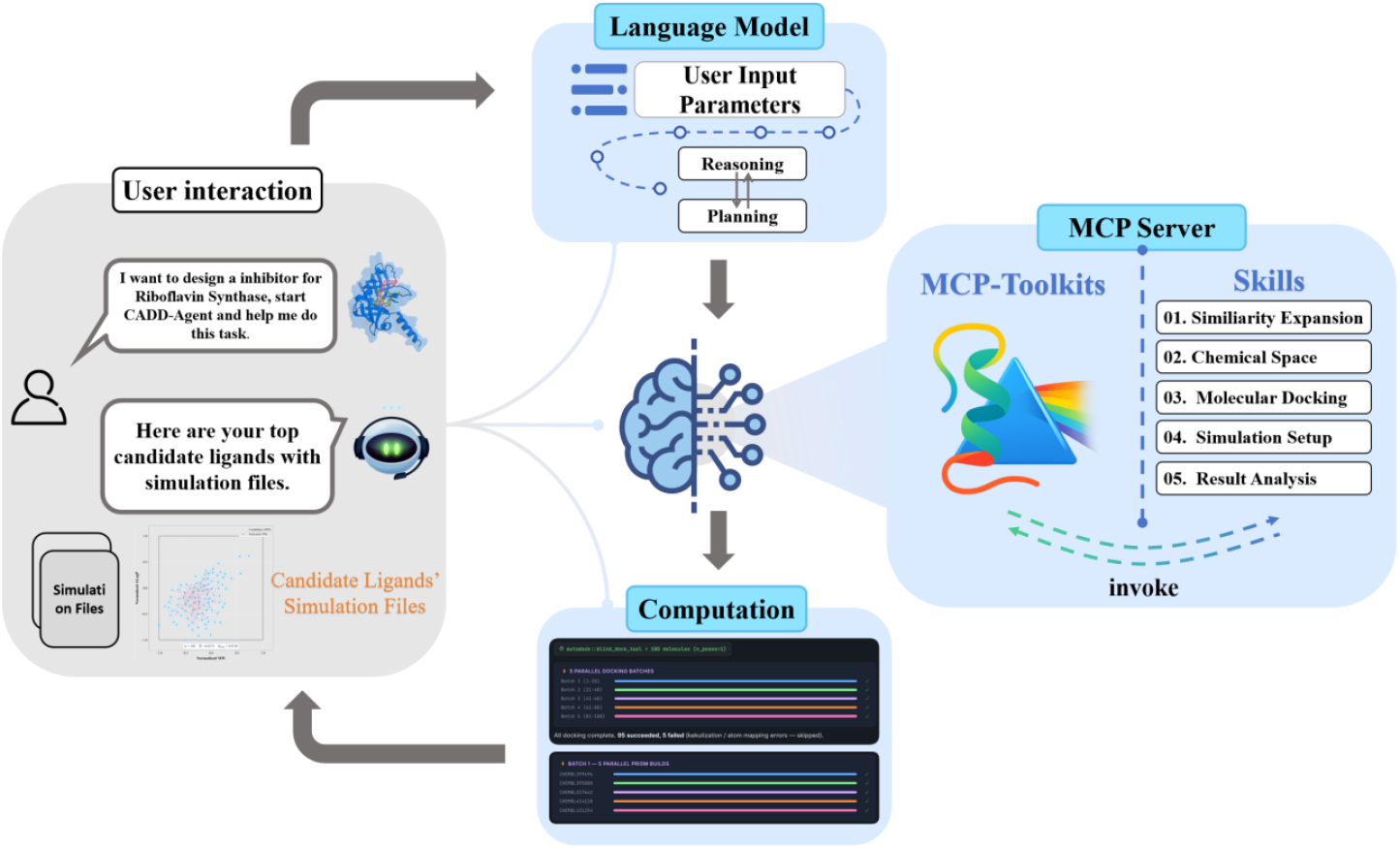
Workflow of the CADD-Agent. A user’s natural-language research intent is interpreted by an LLM guided by a predefined expert workflow (six-stage pipeline prompt encoding domain constraints and quality gates). Computational tools--ChEMBLfind, MolScope, AutoDock Vina MCP, and PRISM--are exposed as independent MCP servers, enabling the LLM to chain tool calls with human-in-the-loop confirmation at each stage.

### 2.3 Simulation Configuration

Simulation parameters are governed by the YAML configuration system (CLI arguments > user config > defaults) that automates .mdp file editing. PRISM automatically generates MDP files for energy minimization, NVT/NPT equilibration, and production MD with validated defaults: velocity-rescale thermostat, C-rescale barostat^24^, PME electrostatics (1.0 nm cutoff, 0.16 nm Fourier spacing), LINCS constraints on hydrogen bonds, and a 500 ns production simulation with 2 fs as time step. Multi-ligand workflows build independent systems in parallel with structured output directories.

### 2.4 Enhanced Sampling: REST2

PRISM automates the setup of Replica Exchange with Solute Tempering 2^25^ (REST2) simulation. The temperature ladder distributes *N* replicas between *T*_*ref*_ (default: 310 K) and *T*_max_ (default: 450 K) using a geometric (exponential) progression:

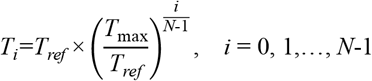

yielding a scaling factor *λ*_*i*_=*T*_*ref*_ / *T*_*i*_ for each replica, where *λ*_0_=1 (unscaled) and *λ*_*N*-1_=*T*_*ref*_/*T*_max_ (maximally scaled). The geometric spacing ensures approximately uniform exchange acceptance ratios between adjacent replicas. The partial tempering scheme scales solute Lennard-Jones well depths by λ, partial charges by 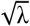, and bonded force constants by λ (intramolecular) or 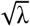 (solute-solvent), preserving solvent-solvent interactions at T_ref_. PRISM generates per-replica topology files, shared equilibration .mdp files, and a single orchestration script for the complete REST2 workflow.

### 2.5 Free Energy Calculation Modules

#### MM/PBSA

PRISM automates endpoint free energy analysis with two backends (gmx_MMPBSA^26^ and AMBER MMPBSA.py via ParmEd^27^ topology conversion) and two modes: single-frame for rapid high-throughput assessment and trajectory-based for conformational averaging. The calculation decomposes binding free energy into van der Waals, electrostatic, polar solvation, and nonpolar solvation contributions.

#### PMF with Automated Pulling Direction Optimization

The PMF module constructs umbrella sampling systems for ligand unbinding free energy profiles. Its key methodological contribution is an automated pulling direction optimization algorithm based on Metropolis-Hastings sampling on the unit sphere *S*^2^ with simulated annealing. The pulling direction 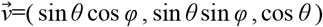 is optimized by minimizing an objective function that quantifies steric hindrance along the candidate pathway.

In the default pocket clearance mode, the binding pocket is defined as all protein heavy atoms 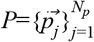 within cutoff distance *r*_c_ (default: 4.0 Å) of the ligand. The objective function uses point-to-line distances between pocket atoms and ligand pulling rays:

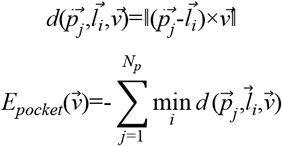

In the whole-protein collision mode, the displacement 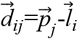 is decomposed into parallel and perpendicular components 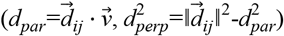, and the collision step range is solved analytically with 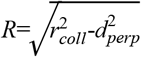 :

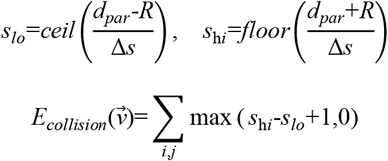

reducing complexity from *O*(*N*_*l*_×*N*_*p*_×*S*) to *O*(*N*_*l*_×*N*_*p*_) per evaluation. The MH sampler proposes directions via Gaussian perturbation *σ*=0.15*rad*, accepts with probability *P*_*accept*_=min (1, exp (-Δ*E*/*T*)), and follows an exponential cooling schedule *T*_*k*_=*T*_0_*α*^*k*^(*T*_0_=2.0,*α*=0.99995). After convergence, the complex is rotated to align the optimal direction with the z-axis, and steered MD and umbrella sampling inputs are generated.

### 2.6 PRISM-FEbuilder

PRISM-FEbuilder automates the construction of ligand perturbation systems for relative binding free energy calculations. A key challenge in FEP setup is establishing atom-wise correspondence between reference and transformed ligands while managing electrostatic differences across the perturbation. FEbuilder addresses this through distance-based atom mapping (default cutoff: 0.6 Å) that classifies atoms into three categories: common (shared between both ligands), transformed (unique to each state), and surrounding (position-matched but with divergent atom types or charges). For common atoms with minor charge differences, the module applies charge assignment strategies including reference-state preservation, mutant-state preservation, or arithmetic averaging to maintain electrostatic consistency across the shared scaffold. Atoms with significant parameter divergence are retained as surrounding atoms, preserving state-specific characteristics. The framework generates single-topology GROMACS .itp files where dual-state parameters are encoded via typeB/chargeB columns, with dummy atom types (DUM_*) handling non-interacting states. λ-window configuration with soft-core potentials completes the perturbation setup for both bound and unbound simulations, enabling standardized conversion of related ligand series into FEP-ready inputs.

### 2.7 Trajectory Analysis and Visualization

PRISM includes an integrated analysis suite built on MDTraj^28^, providing caculations of RMSD, protein-ligand contacts (hysteresis-based, enter: 3.5 Å, exit: 4.0 Å), hydrogen bonds, interatomic distances, solvent-accessible surface area (SASA), dihedral distributions, and conformational clustering. An interactive HTML visualization generator renders contact networks as node-edge graphs with frequency-weighted edges.

To improve the robustness of contact detection against thermal fluctuations, PRISM employs a hysteresis-based definition in which contacts are formed when the distance falls below an entry threshold (3.5 Å) and are only broken when exceeding an exit threshold (4.0 Å). This approach prevents rapid switching artifacts and enables stable characterization of persistent interactions over time, as illustrated in Figure S2-D. Time-resolved analyses further quantify protein-ligand interaction dynamics at multiple levels. PRISM tracks minimum distances between ligand atoms and selected residues, enabling identification of transient versus stable contacts (Figure S2-A), and computes cumulative contact formation to assess the evolution of interaction diversity along the trajectory (Figure S2-B). In addition, the number of maintained (hysteresis-stabilized) contacts is monitored to characterize binding stability (Figure S2-C), while per-residue contact probabilities are calculated to identify key interaction hotspots (Figure S2-F).

A hierarchical plotting system produces publication-quality figures, including time-series traces, violin plots of normalized contact frequencies (Figure S2-E), and ranked residue interaction profiles (Figure S2-F). A built-in multi-system comparison module further enables batch evaluation across compound libraries, supporting consistent analysis in high-throughput screening workflows.

### 2.8 CADD-Agent: Expert-Workflow-Driven AI Interface

As a natural extension of its modular architecture, PRISM serves as the computational backbone of the CADD-Agent, an AI agent that orchestrates multi-step drug discovery workflows. The CADD-Agent separates concerns between a predefined expert workflow — a structured, natural-language protocol that encodes domain knowledge as stage-gated instructions—and a large language model (LLM) that reads this protocol and serves as an adaptive orchestrator. All computational tools, including ChEMBLFind for bioactivity database mining, MolScope for chemical space coverage optimization, AutoDock^29^ Vina for molecular docking, and PRISM for MD system construction and analysis) are exposed as independent MCP servers, allowing the LLM to chain them by passing outputs from one tool as inputs to the next.

The expert workflow encodes decisions that would otherwise require specialist experience: recommended library sizes, force field combinations, parameter constraints (e.g., prohibiting modification of default box size and ionic strength), fault-tolerance rules for docking failures, and quality gates at each stage. The LLM translates high-level research intent into concrete tool calls while maintaining a mandatory human-in-the-loop confirmation pattern before each stage. This architecture ensures scientific rigor through protocol constraints while providing the natural-language flexibility, error recovery, and adaptive orchestration that static scripts cannot offer.

## 3. RESULT

### 3.1 PRISM-CADD-Agent Enabled Hierarchical Screening of Substrate-Analog Inhibitors Against Riboflavin Syntheses

To demonstrate how PRISM and its CADD-Agent orchestration layer enable an end-to-end inhibitor discovery workflow, we conducted a pilot screening study targeting riboflavin synthase (EC 2.5.1.9, PDB: 1KZL)^30^. Riboflavin synthase catalyzes the final step in the riboflavin (vitamin B2) biosynthesis pathway—the dismutation of two molecules of 6,7-dimethyl-8-ribityllumazine into riboflavin and pyrimidinedione. Because this pathway is essential in many bacterial species yet entirely absent in mammals, riboflavin biosynthetic enzymes have been recognized as attractive antimicrobial targets with inherent selectivity. Structurally, the enzyme functions as a homotrimer: each ∼23 kDa monomer contains N- and C-terminal beta-barrel domains connected by a linker, and the catalytic active sites form exclusively at the interface between adjacent monomers, with a C-terminal alpha-helix playing a critical role in maintaining the trimeric assembly.

The entire screening workflow was orchestrated by the CADD-Agent operating under PRISM’s expert workflow protocol (Figure 5A). ChEMBLFind was first used to query the ChEMBL database^31^ for bioactive molecules associated with riboflavin synthase and its substrate analogs, yielding an initial candidate pool of 903 compounds. To ensure that downstream docking covered a chemically diverse and representative subset, MolScope was applied to optimize chemical space coverage in the normalized MW ⊗ ALogP descriptor space. From the 903 candidates, 100 representative molecules were selected using maximin coverage optimization (coverage radius R = 0.0397, minimum inter-point distance *d*_min_=0.0794), ensuring uniform sampling across the accessible chemical space (Figure 5B). These 100 representatives were then subjected to blind docking against 1KZL using AutoDock Vina. The top 10 compounds ranked by docking score were carried forward to the physics-based assessment stage.

**Figure 5.**
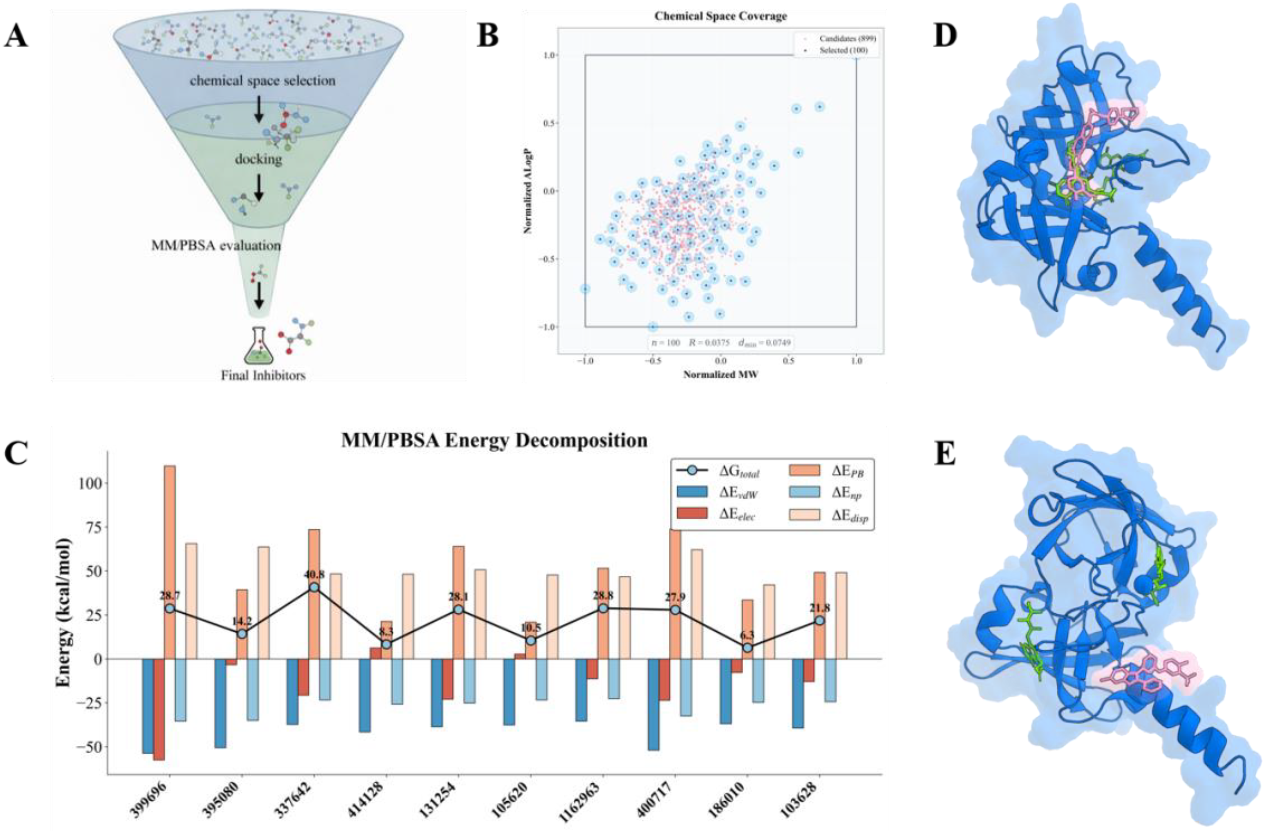
Hierarchical screening of Riboflavin Synthase inhibitors. **(A)** Multi-tier screening funnel: ChEMBL candidate search → chemical space selection → docking → MM/PBSA evaluation. **(B)** Chemical space coverage: 100 representatives selected from 903 compounds via maximin optimization in normalized MW⊗ALogP space. **(C)** MM/PBSA energy decomposition for the top-10 docking-ranked candidates, with Δ*G*_*total*_ ranging from 6.5 to 40.8 kcal/mol. **(D)** A top-5 candidate (pink) overlaps with the co-crystallized substrate analog (green) at the active site, validating the pipeline. **(E)** The top-ranked candidate (CHEMBL186010) binds at the C-terminal α-helix base critical for homotrimerization, suggesting allosteric inhibition via oligomeric interface disruption.

For each of the 10 docking-prioritized candidates, PRISM automatically constructed explicit-solvent protein-ligand systems (AMBER14SB protein force field, GAFF2 ligand parameters, Gaussian HF/6-31G* RESP charges, PROPKA-based protonation) and performed single-frame MM/PBSA binding free energy calculations. The energy decomposition analysis (Figure 5C) revealed that all 10 candidates exhibited favorable van der Waals (Δ*E*_*vdW*_) and electrostatic (Δ*E*_*elec*_) contributions, but these were partially offset by the polar solvation desolvation penalty (Δ*E*_*PB*_) and dispersion corrections (Δ*E*_*disp*_). The resulting Δ*G*_*total*_ values ranged from 6.5 to 40.8 kcal/mol, with the five most favorable candidates being CHEMBL186010 (6.5 kcal/mol), CHEMBL414128 (8.3 kcal/mol), CHEMBL105620 (10.5 kcal/mol), CHEMBL395080 (14.2 kcal/mol), and CHEMBL103628 (21.8 kcal/mol).

Structural inspection of the top-5 binding poses yielded two noteworthy findings. First, one of the top-5 candidates bound precisely at the crystallographically resolved substrate binding site, with its docked pose closely overlapping the position of the co-crystallized ligand 6-carboxyethyl-7-oxo-8-ribityllumazine (Figure 5D, green: crystal ligand; pink: docked candidate). This concordance between computational prediction and experimental structure validates the reliability of the screening pipeline. Second, and more intriguingly, the top-ranked candidate (CHEMBL186010, Δ*G*_*total*_=6.5*kcal*/*mol*) did not bind at the canonical active site. Instead, it occupied a pocket located at the base of the C-terminal alpha-helix (Figure 5E), a structural element that is indispensable for homotrimerization. Since riboflavin synthase is obligately trimeric—with active sites forming exclusively at the inter-subunit interface—ligand binding at this site could potentially disrupt trimerization and abolish catalytic activity through an allosteric mechanism. This strategy of targeting oligomeric interfaces, rather than the catalytic pocket itself, has well-established precedents in drug design, including the disruption of the TNF-*α* trimer by SPD304 and the modulation of ClpP protease oligomerization by acyldepsipeptide antibiotics. Our result suggests that riboflavin synthase inhibition may be achievable not only through competitive substrate analogs but also through trimerization-disrupting compounds, offering a complementary and potentially resistance-resistant inhibition strategy.

### 3.2 Benchmark of PRISM-FEbuilder

To evaluate the performance of the FEbuilder module for relative binding free energy calculations, we benchmarked three representative protein-ligand systems: HIF-2α, T4 lysozyme L99A, and p38α kinase, all with experimentally measured ligand affinities available (Fig. 6). The HIF-2α and p38α series were selected from established protein-ligand benchmark collections developed for rigorous binding affinity evaluation, whereas T4-lysozyme L99A was included as a classical model system featuring a buried hydrophobic cavity and chemically diverse ligands. For each target, ligand perturbation cycles were constructed on the basis of structural similarity, and relative free energy perturbation calculations were performed along the resulting transformation networks. Ligands were parameterized using MATCH, and the benchmark was carried out in a CHARMM-compatible setup with multiple repeated runs for each perturbation. We employed standard alchemical equilibration and production settings, including explicit-solvent bound and unbound simulations, PME electrostatics, a 12.0 Å nonbonded cutoff, a 1.0 fs timestep, and 310 K NPT equilibration prior to production FEP calculations.

**Figure 6.**
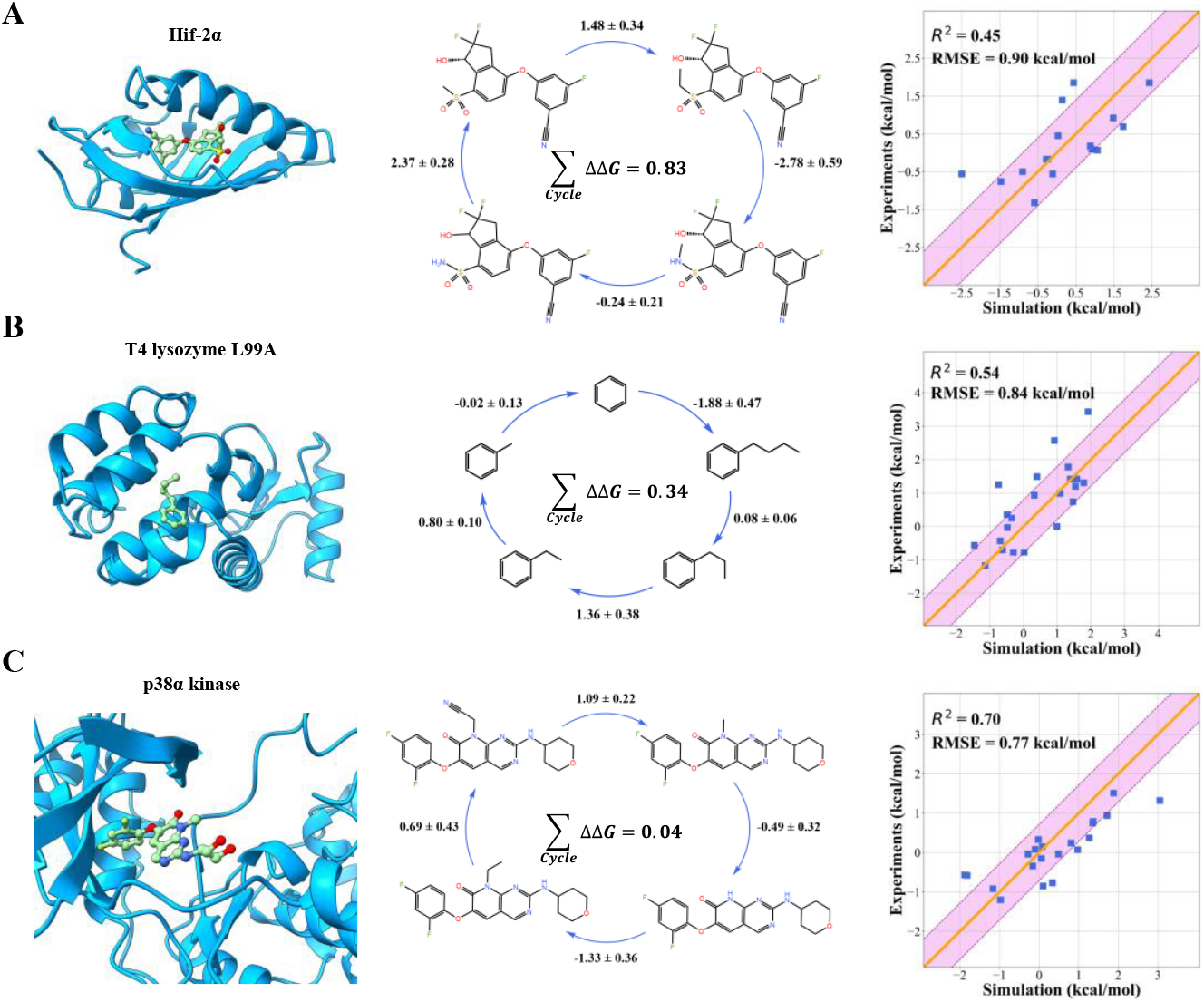
Benchmark of PRISM-FEbuilder. Relative binding free energy calculations for **(A)** HIF-2α, **(B)** T4-lysozyme L99A, and **(C)** p38α kinase. Left: protein structures; center: alchemical perturbation networks with cycle-closure constraints; right: calculated vs. experimental ΔΔ*G* scatter plots (yellow line: equation *x* = *y*; pink band: ±1 std). RMSE values are 0.90, 0.72, and 0.77 kcal/mol with R^2^of 0.45, 0.54, and 0.70, respectively.

Across all three systems, the perturbation cycles were generally well behaved, with small cycle-closure hysteresis, indicating internally consistent free energy networks. As shown in Fig. 6, the calculated relative affinities agreed well with experiment, yielding RMSE values of 0.90 kcal/mol for HIF-2α, 0.72 kcal/mol for T4-lysozyme L99A, and 0.77 kcal/mol for p38α kinase, with corresponding R^2^ values of 0.45, 0.54, and 0.70, respectively. Most perturbations showed unsigned errors within approximately 1 kcal/mol. These results support the robustness and practical reliability of systems generated by FEP module for relative free energy calculations across chemically related ligand series.

## 4. DISCUSSION

In summary, PRISM provides an integrated and high-throughput simulation infrastructure that enables scalable, hierarchical protein-ligand screening workflows for CADD agents. Rather than functioning as a collection of isolated scripts, PRISM unifies ligand parameterization, system construction, simulation setup, enhanced sampling, multi-tier binding free energy calculation, and trajectory analysis within a single framework. In doing so, it enables a hierarchical screening strategy that combines docking for rapid filtering, molecular dynamics for stability assessment, PMF for dissociation thermodynamics, and FEP for fine-grained discrimination. The riboflavin synthase case study demonstrates the end-to-end capability of this framework, from candidate library assembly to binding-pocket characterization, and further highlights its potential to support biologically meaningful hypothesis generation, including the identification of a putative allosteric inhibition site at the oligomerization interface.

At the same time, the performance of any computational screening pipeline remains dependent on input quality and modeling choices, including protein structure accuracy, ligand parameterization, protonation state assignment, sampling sufficiency, and force-field reliability. Although PRISM automates these procedures and provides validated default settings, it cannot guarantee that default choices are optimal for every system, particularly for targets with unusual chemistry, pronounced conformational heterogeneity, or poorly characterized binding sites. In addition, the hierarchical screening strategy necessarily involves trade-offs: compounds excluded at early stages may include false negatives that would otherwise survive more rigorous downstream evaluation. Finally, because the present study focuses on a single target system, broader benchmarking across diverse protein families, binding environments, and ligand chemotypes will be important for establishing the generalizability of both the platform and the associated screening paradigm. Taken together, PRISM provides a practical foundation for scalable, agent-enabled simulation workflows in computer-aided drug design.

## Supporting information

cover letter

## Data Available

The source code of **PRISM** is now available at both Github and Zenodo:

**Github:** https://github.com/AIB001/PRISM

**Zenodo:** https://zenodo.org/records/19163575

Plugins for **PRISM-CADD-Agent** are available at:

**ChEMBLfind**: https://github.com/AIB001/chemblfind

**MolScope**: https://github.com/AIB001/molscope

**Autodock Vina MCP**: https://github.com/AIB001/AutodockVina_MCP

## Acknowledgement

We thank Yue Wu from Department of Chemistry, University of Wisconsin-Madison, Lianxue Zhang from College of Life Sciences, Zhejiang University for helpful discussions. This work was partially supported by the National Key R&D Program of China (2024YFA1306400, 2021YFA1201200, 2024YFA1307500), the National Natural Science Foundation of China (U1967217), the National Center of Technology Innovation for Biopharmaceuticals (NCTIB2022HS02010), Shanghai Artificial Intelligence Lab (P22KN00272), the National Independent Innovation Demonstration Zone Shanghai Zhangjiang Major Projects (ZJZX2020014), the Starry Night Science Fund of Zhejiang University Shanghai Institute for Advanced Study (SN-ZJU-SIAS-003), Zhejiang University Global Partnership Fund (188170+194452505), and the Sichuan Science and Technology Program (2025YFHZ0065).

## Supplementary information

**Figure S1.**
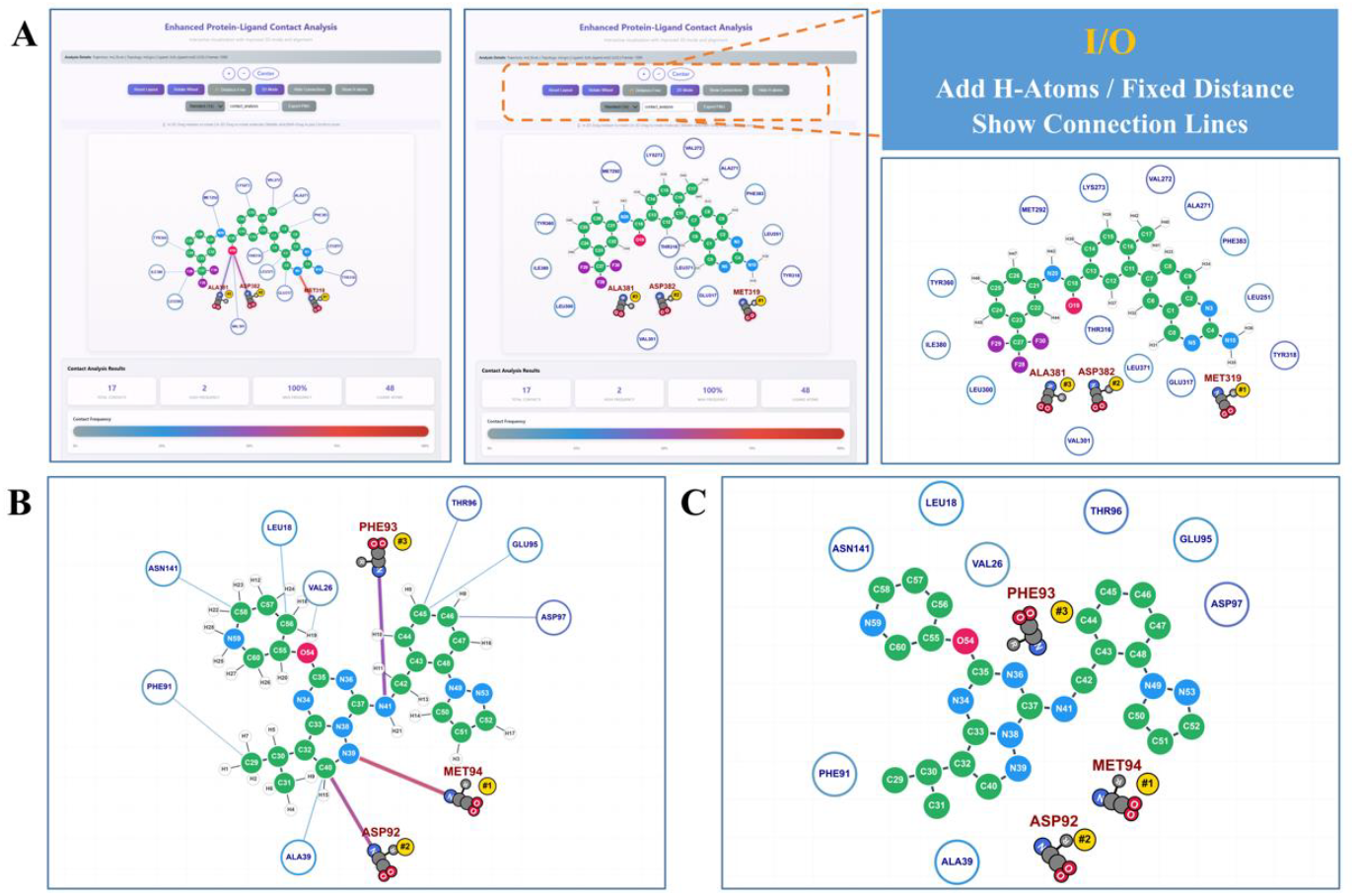
Interactive HTML visualization of protein–ligand contact networks. **(A)** The web-based interface renders ligand and surrounding residues as node–edge graphs, with toggleable options for hydrogen atom display, fixed-distance layout, and connection line visibility (right inset). **(B–C)** Two representative contact network views showing frequency-weighted edges between ligand atoms and contacting residues (e.g., PHE93, MET94, ASP92), with node color encoding residue type and edge thickness reflecting contact frequency.

**Figure S2.**
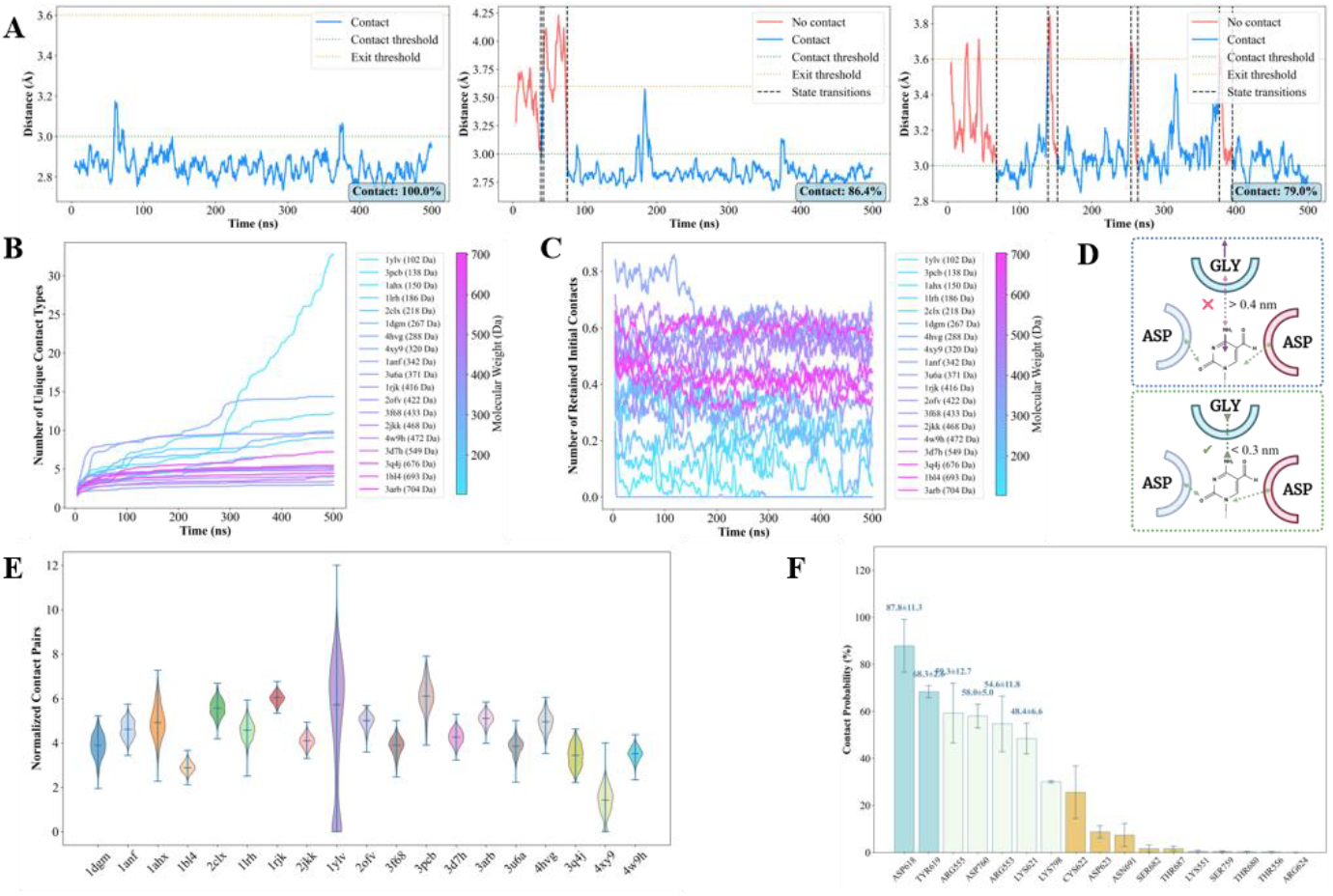
Trajectory post-analysis outputs generated by PRISM. **(A)** Time-resolved protein–ligand minimum distance traces for three representative residue–ligand pairs, with contact and exit thresholds (enter: 3.5 Å, exit: 4.0 Å) indicated by dashed lines; state transitions between contact and non-contact are marked. Overall contact percentages are annotated. **(B)** Cumulative number of unique contact types over time, stratified by residue. **(C)** Number of maintained (histered) contacts over time for individual residues, reflecting binding stability. **(D)** Schematic of the hysteresis-based contact definition: contacts form when distance falls below 0.3 nm and break only when distance exceeds 0.4 nm, preventing rapid state switching.**(E)** Violin plots of normalized contact pairs per residue across the trajectory. **(F)** Per-residue contact probability (%) ranked by frequency, identifying the most persistent interaction partners.

**Figure S3.**
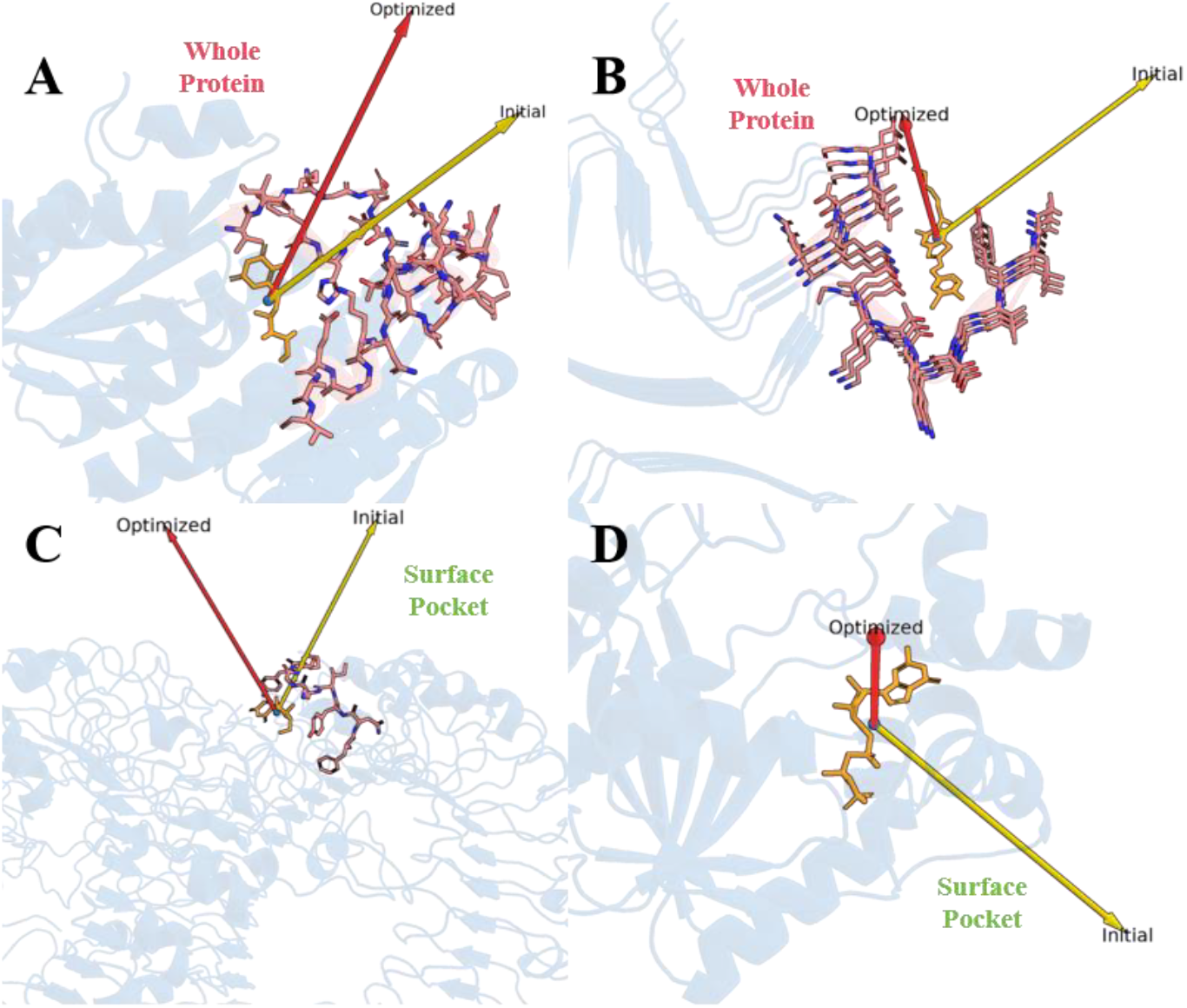
Comparison of pulling direction optimization modes for PMF setup. Yellow arrows indicate the naive centroid-to-centroid vector between protein and ligand; red arrows show the optimized pulling direction from PRISM. **Upper panels:** Whole-protein collision mode applied to two systems with deeply buried ligands, where the naive direction passes through the protein interior while the optimized direction identifies a sterically unobstructed exit pathway. **Lower panels:** Surface pocket mode applied to surface-bound ligands, where the naive direction similarly traverses neighboring structural elements, whereas the optimized direction avoids steric collisions with the pocket walls. Both modes demonstrate clear improvement over the conventional approach.

**Figure S4.**
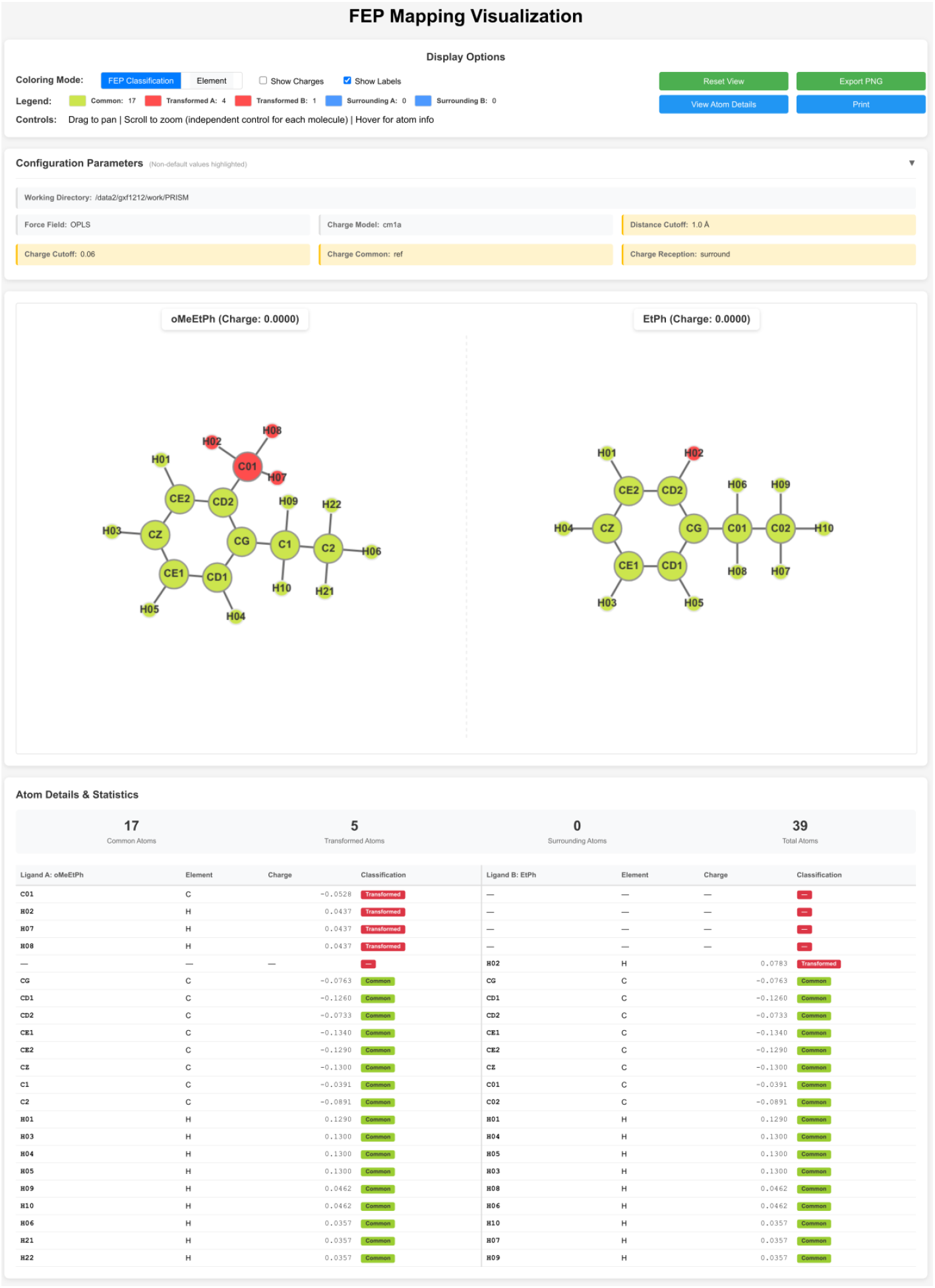
Interactive HTML visualization of the hybrid topology generated by PRISM-FEbuilder for alchemical transformation between o-methoxyethylphenyl (oMeEtPh) and ethylphenyl (EtPh) ligands using OPLS force field. The web interface displays side-by-side 2D molecular structures with color-coded atom classification based on distance-based mapping: common atoms (green, shared scaffold), transformed atoms (red), and surrounding atoms (blue). A comprehensive correspondence table lists atom names, elements, partial charges, and FEP classifications for both ligand states, facilitating verification of dual-state parameter encoding in GROMACS .itp files via typeB/chargeB columns. Users can independently rotate, zoom, and pan each molecular view, with hover tooltips revealing detailed atom information. This visualization aids in validating the automated hybrid topology construction and charge assignment strategies (reference-state, mutant-state, or arithmetic averaging) that underpin accurate relative binding free energy predictions.

**Table S1.**
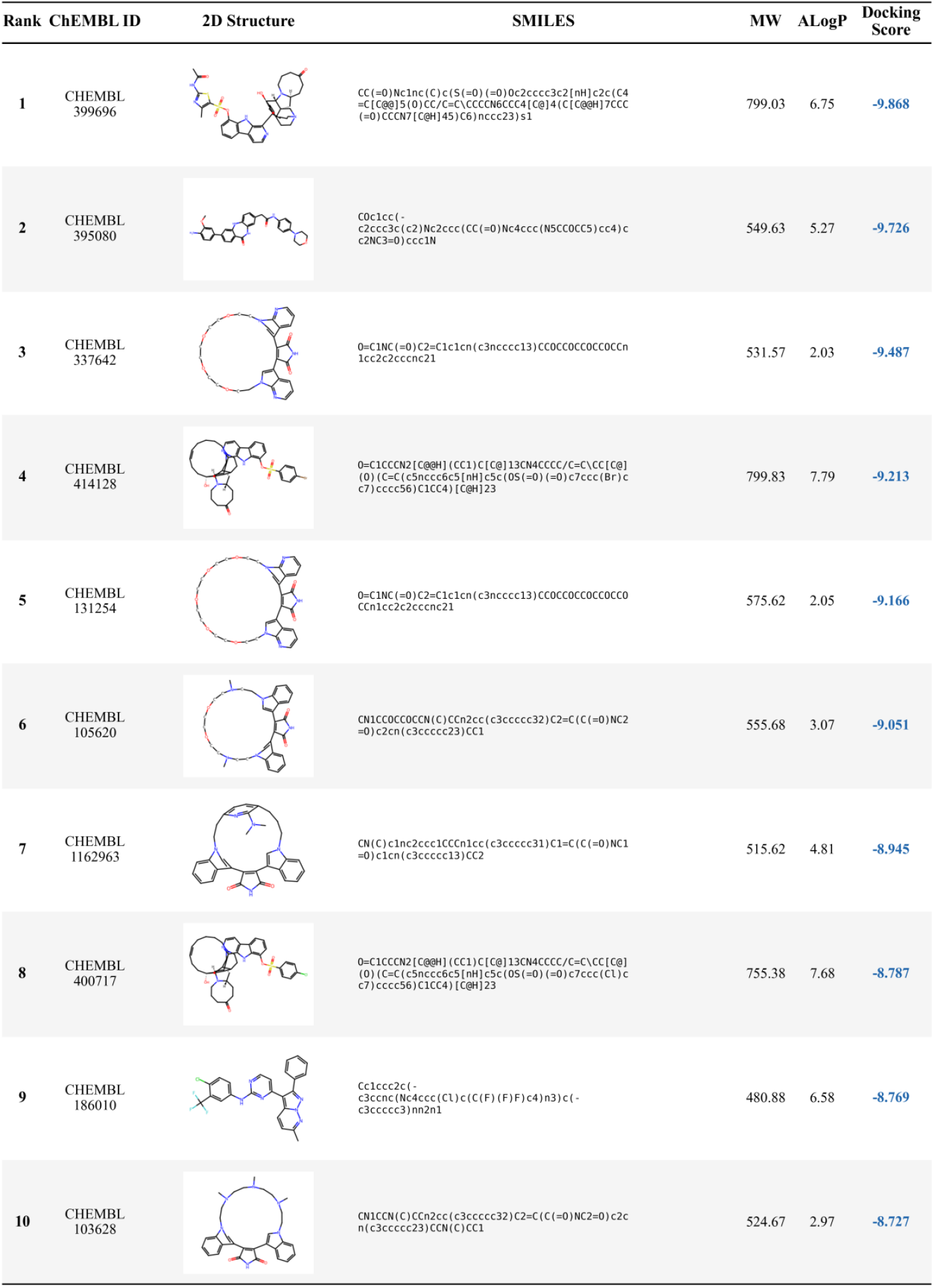
Top-10 docking-ranked candidates from the hierarchical screening against riboflavin synthase (PDB: 1KZL). Compounds were selected from 100 chemical space representatives via blind docking with AutoDock Vina. Listed properties include 2D structure, SMILES notation, molecular weight (MW), calculated lipophilicity (ALogP), and docking score (kcal/mol).

