## Supplementary material for "PRISM: A High-Throughput Simulation Infrastructure for CADD Agents": cover letter

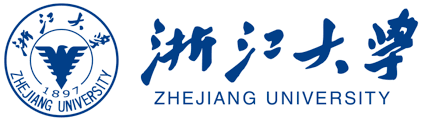
Qiushi Chair Professor and Director
Institute of Quantitative Biology
Zhejiang University, China

April 2, 2026

Dear editors,

We are pleased to submit the manuscript entitled **"PRISM: A High-Throughput Simulation Infrastructure for CADD Agents"** for your kind consideration of publication. The following is a brief description of our work and its significance for computer-aided drug design.

Modern computer-aided drug design increasingly relies on molecular simulation for candidate evaluation, mechanistic interpretation, and binding free energy estimation. However, in practice, protein–ligand simulation workflows remain highly fragmented across multiple tools, making reproducible, scalable, and iterative screening difficult. We developed PRISM to address this bottleneck by providing an integrated, high-throughput simulation infrastructure that unifies ligand parameterization, automated system construction, enhanced sampling, multi-tier free energy evaluation, and trajectory analysis within a single GROMACS-based Python platform. Through the Model Context Protocol (MCP), PRISM serves as the computational backend for CADD-Agent, enabling large-language-model-based orchestration of hierarchical drug screening workflows while preserving expert-defined constraints and reproducibility. In this way, our work bridges rigorous molecular simulation with AI-driven scientific workflow automation.

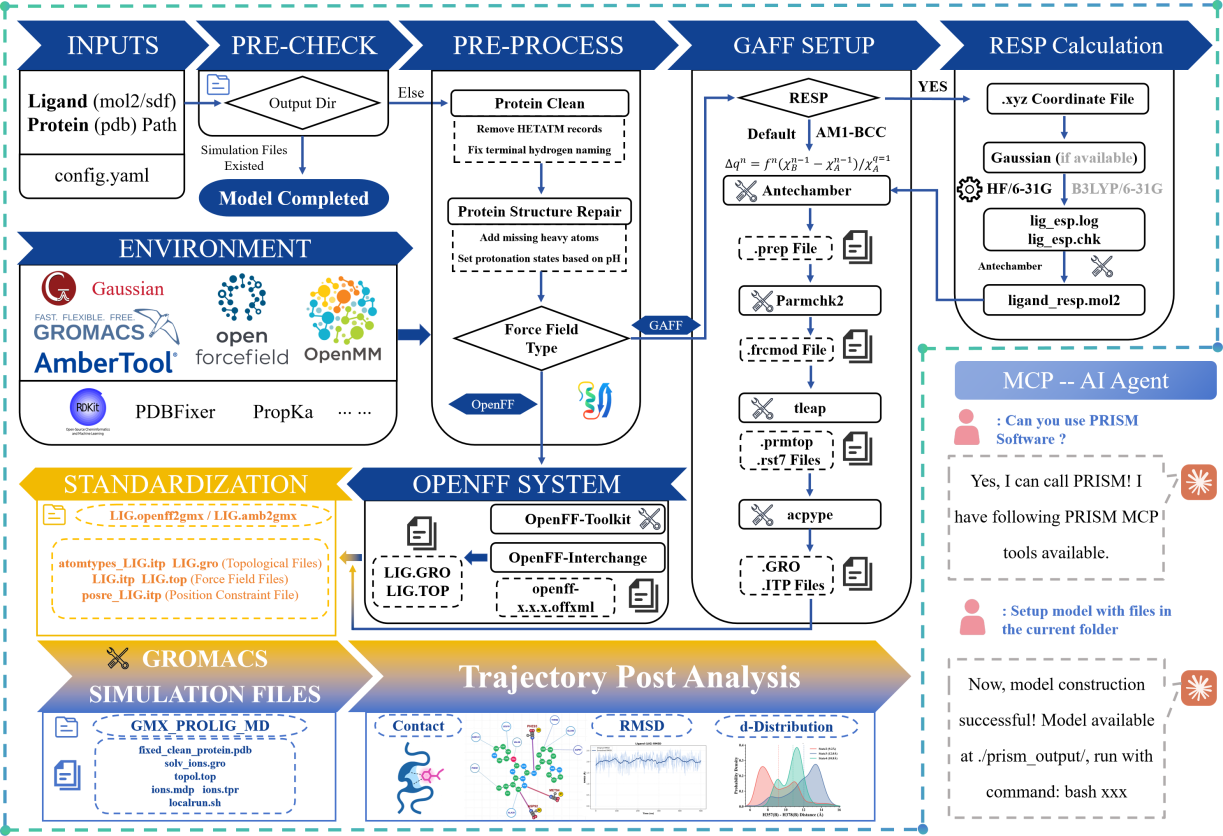

**Figure 1. Architecture of PRISM.**

In summary, this work will be of interest to readers in molecular simulation, computational chemistry, and computer-aided drug design, particularly those interested in scalable simulation workflows and AI-assisted molecular discovery. PRISM provides not only a practical platform for high-throughput protein–ligand simulation, but also a foundation for future agent-enabled CADD applications.

Thank you for your consideration. We would be grateful for the opportunity to have our work evaluated for publication.

Thanks.

Sincerely,

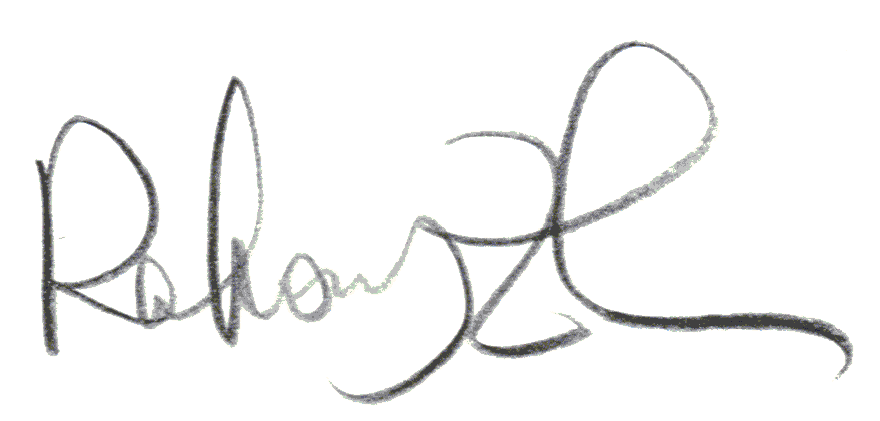

Ruhong Zhou, PhD

Zhejiang University and Columbia University
